# Sexual dimorphism in head size in wild burying beetles (*Nicrophorus vespilloides*)

**DOI:** 10.1101/2023.11.01.565177

**Authors:** Jack M.L. Smith, Andrew M. Catherall-Ostler, Rahia Mashoodh, Rebecca M. Kilner

**Author notes:** Correspondence: Jack M. Smith, Department of Zoology, University of Cambridge, Downing Street, Cambridge CB2 3EJ.

## Abstract

1. Male burying beetles (*Nicrophorus vespilloides*) carry an additional abdominal segment and produce pheromones but otherwise the species is thought to be sexually monomorphic. Both sexes bear bright orange bands on their black elytra, which probably function as part of a warning display rather than in mate choice.
2. Although larger individuals are more likely to win contests to secure the small carrion required for reproduction, both sexes compete for this vital breeding resource. In wild populations, the sexes are alike in the mean and distribution in their body size.
3. Here we describe a form of sexual size dimorphism in wild populations that has previously been overlooked. We show that males have wider heads than females, for any given body size, and that the scaling relationship with body size is hyperallometric in males, but isometric in females. We also show how absolute head width, and the extent of sexual dimorphism in head width differs among wild populations which are within c.10km of each other.
4. We suggest that head size dimorphism could be adaptive, and could be due to divergent selection arising from task specialisation during biparental care, since the duties of care favoured by males are likely to require a greater bite force.

## Introduction

Instances of sexual dimorphism in animals are usually attributed to sexual selection, which arises either during competition for mating opportunities or through mate choice (Clutton-Brock, 2009, 2007). However, these two contexts do not fully capture all the ways in which males and females behave differently during reproduction (Clutton-Brock, 2009). Consequently, there are additional sources of selection at work which can further explain why the sexes might differ in their morphology (Clutton-Brock, 2009; Shine, 1989; Tosto et al., 2023). For example, when parents cooperate to raise offspring together, the various duties of care are frequently divided up differently between the sexes. The resulting task specialisation potentially imposes selection for morphologies that diverge adaptively according to the tasks that each sex then undertakes (Hedrick and Temeles, 1989; Shine, 1989; Slatkin, 1984). One sex might possess a brood patch to warm the clutch more effectively, for instance, as is typical in bird species where incubation is confined to a single sex (Höhn and Cheng, 1965). Or one sex might possess larger mandibles, as described in female megachilid bees, who apparently use these structures to construct a nesting burrow (Shine, 1989). Selection could act differently on each sex during foraging too, either to prevent competition between the sexes for limited resources or to enhance the foraging efficiency of a pair provisioning offspring (Shine, 1989).

When multiple sources of selection act divergently on males and females at the same time, it can be hard to link divergence in specific traits to specific sources of selection (Clutton-Brock, 2009; Shine, 1989). Our goal here is describe and quantify the extent of divergence between male and female burying beetles *Nicrophorus vespilloides* in a trait that is unlikely to be explained by conventional accounts of sexual selection.

*Nicrophorus* spp are typically described as sexually monomorphic (Al Shareefi and Cotter, 2019; Gibbs et al., 2008; Sikes et al., 2002). Indeed, in *N. vespilloides* the sexes are known to differ in only two ways morphologically. Males possess an extra abdominal segment (Easton, 1979) and are capable of producing pheromones to attract females (Müller and Eggert, 1987). Males are colourful, bearing orange patterning on their black elytra, but so too are females and to the same degree (Lindstedt et al., 2017). The identical bright colouration of both sexes is more likely to be part of a warning display than to function in mate choice (Lindstedt et al., 2017). Body size, and the extent to which it varies, do not differ between the sexes either (Jarrett et al., 2017; Sun et al., 2020).

The unusual natural history of the burying beetle seemingly accounts for the lack of sexual selection, and the corresponding lack of divergence in morphology. Reproduction in *N. vespilloides* centres on the dead body of a small vertebrate, such as a rodent or songbird. Each sex can locate the volatiles emitted by a decomposing body (Trumbo et al., 2021) and can fly for several kilometres to locate carrion to breed upon (Attisano and Kilner, 2015). Together, a pair converts the dead body into a nest for their larvae. They strip off the fur or feathers, roll the flesh into a ball and inter it in a shallow grave, all the while smearing the meat with fluids from their mouth and anus. While they are burying the body, the female lays her eggs in the surrounding soil. The larvae hatch and crawl to the balled carcass, which becomes their edible nest. There they feed themselves on the flesh (Smiseth et al., 2003) and are tended by both parents. Parents guard their larvae and nest from infanticidal takeover by rival *Nicrophorus* spp, maintain the carrion nest, and transfer fluids to larvae by oral trophallaxis. Roughly a week later, the larvae crawl away from the scant remains of the carcass to pupate in the soil and each parent flies off independently in search of new breeding opportunities.

The burying beetle’s natural history, and its dependence on carrion, has three important consequences for the way that selection acts on each sex. First, it greatly limits the scope for mate choice, and reduces the scale of any benefits that it might bring. Beetles are solitary except when they gather at a dead body to feed or breed (Scott, 1998), or when males attract females through pheromonal signalling (Müller and Eggert, 1987). They apparently mate indiscriminately when given the opportunity (House et al., 2008; Mattey and Smiseth, 2015) and females can arrive at a dead body having already been inseminated by multiple males (Müller et al., 2007). Therefore, there is no compelling evidence for mate choice by either sex. Second, the beetles’ dependence on a rare bonanza resource to breed means that competition for carrion within each sex is anyway likely to be a far more significant component of sexual selection than mate choice (see Clutton-Brock, 2009; Royle and Hopwood, 2017). Yet *both* sexes are exposed to this selection pressure. Competition ensues if more than one member of each sex happens upon the same dead body and successful competitors tend to be larger (Royle and Hopwood, 2017). Although the extent of competition for carrion can influence body size in wild populations, there is no evidence that selection on body size differs between the sexes (Sun et al., 2020).

The beetles’ capacity to carry out biparental care introduces a different context in which selection can potentially act divergently on males and females. In *N. vespilloides*, each sex can potentially carry out all the duties of care. However, when parents care for offspring together, males invest relatively more effort in making the carrion nest (De Gasperin et al., 2016; Jarrett et al., 2023) and defending the brood and nest from infanticidal takeover (Müller et al., 1998; Scott, 1998; Trumbo, 2007, 1994), whereas females put relatively more effort into offspring provisioning and post-hatching carrion maintenance (Smiseth et al., 2005; Walling et al., 2008). Task specialisation during care could cause selection to act differently on each sex, to yield traits that are sexually dimorphic. Here we provide evidence for sexual dimorphism that has previously been overlooked in *N. vespilloides*, in a trait that might be linked to task specialisation during care.

Our preliminary observations of laboratory populations of *N. vespilloides* indicated that males have larger heads for their body size than females (Smith, 2023). Here we describe in detail how wild populations of *N. vespilloides* are sexually dimorphic in head size. We 1) report the allometric scaling relationship between head size and body size for each sex; and 2) describe how the extent of sexual dimorphism in head size varies between neighbouring woodland populations.

## Materials & Methods

### Field sites

We studied beetles trapped in seven woods (see Supplementary / Figure 1) at the intersection of Cambridgeshire, Bedfordshire and Huntingdonshire (UK): Buff Wood (Cambridgeshire, N = 37) Cockayne Hatley Wood (Bedfordshire, N = 72), Gamlingay Wood (Cambridgeshire, N = 481), Hayley Wood (Cambridgeshire, N = 283), Potton Wood (Bedfordshire, N = 369), Waresley Wood (Huntingdonshire, N = 282, and Weaveley Wood (Huntingdonshire, N = 204). All are Sites of Special Scientific Interest apart from Cockayne Hayley, which is a County Wildlife Site. Gamlingay, Hayley and Waresley (Bedfordshire, Cambridgeshire & Northamptonshire Wildlife Trust (BCN WT)) and Potton Wood (Forestry Commission) are nature reserves open to the public; Cockayne Hatley Wood and Weaveley Wood are privately owned with no public access, whilst Buff Wood is privately owned but managed by the BCN WT with permit-only access. All are listed in the Ancient Woodlands Inventory for England and Wales. All except Potton Wood are mentioned in the Domesday Book (further details of these woods are given in Catherall-Ostler, 2023).

### Trapping

Burying beetles were caught using the method described in Sun (2020). Japanese beetle traps were hung from the branch of a tree using a piece of string, suspended 1-2 m away from the ground. Each trap consisted of a funnel, the top of which is covered by a baffle - two bisecting plastic dividers that increased beetle capture rate by deflecting beetles flying over the trap into the funnel. The funnel was screwed to a cylinder, and the cylinder was lined with small holes to allow the drainage of excess rainwater. Beetles were prevented from escaping due to the small size of the funnel opening, the baffle and the steep slope of the funnel surface (Fleming et al., 1940). Each was filled to ¾ of its total volume with moistened MiracleGro All-Purpose Compost and a freshly thawed mouse (usually > 20 g) was placed on top of the soil.

All traps were placed in a similar microhabitat (hung off a tree branch ∼2 m high under a closed canopy); traps were left for roughly two weeks before emptying and rebaiting. This period was chosen because a shorter timeframe would have resulted in fewer beetles per trap, whilst a longer interval would have resulted in substantial within-trap deaths (Sun, 2020). The mean length of time between trapping sessions was 14.8 days ([14.4, 15.2] 95% CI). In 2019, trapping began in late March to ascertain when burying beetles emerged from overwintering: no beetles were recorded till the 18^th^ May trapping session. The data reported here were collected in 2021. That year, trapping began in May and ended in October when the numbers caught approached zero.

### Measuring

Each trap was emptied into a separate plastic box (17 x 12 x 6 cm) and brought back to the Department of Zoology, University of Cambridge. Each individual was anaesthetised with carbon dioxide prior to measuring and sexed using Easton’s (1979) observation that males possess an extra abdominal segment.

We used the standard practice of quantifying body size by measuring pronotum width (Jarrett et al., 2017). Pronotum width and head width were measured using Mitutoyo Digital Vernier callipers (resolution = 0.01 mm). We defined pronotum width as the distance between the two widest points of the pronotum, and head width as the distance between the two widest points of the head capsule (see Supplementary / Figure 2).

### Statistical analyses

Analyses were performed using R version 4.1.3.

#### Allometric analyses

To measure the allometric scaling relationship between pronutum width and head width we used both standardised major axis regression (SMA; Smith, 2009; Warton et al., 2006) and ordinary least squares regression (OLS; Egset et al., 2012; Kilmer and Rodríguez, 2017; Pélabon et al., 2014) using the R package ‘*smatr*’. The approach of OLS is to assume that the predictor variable is measured without error (Kilmer and Rodríguez, 2017). This means OLS has a tendency to underestimate slopes when there is measurement error present (Kilmer and Rodríguez, 2017), so we also utilised SMA in our analyses which allows for measurement error in both axes (Green, 1999; Laws, 2003). Allometry is defined as y = αx^β^, where y is the size of the trait of interest, x is body size, α is the allometric intercept, and β is the allometric scaling parameter. Morphological variables were log_10_-transformed prior to allometric analysis. Taking the logarithm of trait size and body size yields a linear relationship, log(y) = log(α) + β log(x), where log(α) is the intercept and β is the slope of the line. For traits that scale perfectly with body size (isometry), β = 1. We compared the scaling (allometric) relationship between head width and pronotum width for males and females: 1) using the entire dataset (all the woodlands pooled), and 2) with separate analyses for each wood. To determine if woods differed in allometric intercept (α, elevation) and allometric scaling (β, slope) between pronotum width and head width, we conducted separate pairwise post-hoc analyses for males and females. For OLS analyses we used Tukey’s correction using the R package ‘*emmeans*’ (Lenth et al., 2019), and for SMA analyses we used Sidak’s correction with the *sma* function in the R package ‘*smatr*’ (Warton et al., 2012).

#### Variation in the extent of dimorphism

In a final analysis, we investigated whether the extent of sexual dimorphism in head width (i.e., the difference between male and female head width) differed among woods, using an ANCOVA with head size as the response variable. Sex and woodland, and their interaction (sex x woodland) were included as explanatory variables, and pronotum width was included as a covariate. Pronotum width was included as a covariate because there is no sexual dimorphism in this trait (Jarrett et al., 2017; Sun et al., 2020) - we also confirmed this in our own analyses using an ANCOVA to model pronotum size as a function of wood and sex (see Supplementary / Figure 3 and Table 1). Finally, if a significant interaction was found, pairwise post-hoc analyses using Tukey’s correction were conducted to test for differences in the degree of sexual dimorphism between woods using the R package ‘*emmeans*’ (Lenth et al., 2019).

**Table 1.** The allometric slopes (β) of the relationship between head width and pronotum width in male and female *Nicrophorus vespilloides* for the entire dataset (‘*woodlands pooled*’), and for each woodland independently. Summary statistics for both ordinary least squares (OLS) and standardised major axis (SMA) regression estimates are shown, p-values indicate testing for a significant deviation from a slope of one (β_1_)

| Woodland | Sex | OLS |  |  |  | SMA |  |  |  |
| --- | --- | --- | --- | --- | --- | --- | --- | --- | --- |
| | | Slope ($\beta$) | CI | p | Allometry | Slope ($\beta$) | CI | p | Allometry |
| Woodlands pooled | M | 1.35 | 1.32, 1.37 | <0.001 | + | 1.41 | 1.39, 1.44 | <0.001 | + |
|  | F | 0.95 | 0.93, 0.98 | <0.001 | - | 1.01 | 0.99, 1.04 | 0.27 | 0 |
| Buff | M | 1.44 | 1.24, 1.64 | <0.001 | + | 1.49 | 1.31, 1.70 | <0.001 | + |
|  | F | 0.44 | -0.07, 0.95 | 0.03 | - | 1.03 | 0.64, 1.66 | 0.91 | 0 |
| Cockayne Hatley | M | 1.23 | 1.06, 1.39 | 0.01 | + | 1.27 | 1.11, 1.45 | 0.002 | + |
|  | F | 0.87 | 0.78, 0.97 | 0.009 | - | 0.94 | 0.85, 1.04 | 0.20 | 0 |
| Gamlingay | M | 1.31 | 1.27, 1.36 | <0.001 | + | 1.37 | 1.32, 1.41 | <0.001 | + |
|  | F | 0.99 | 0.95, 1.02 | 0.47 | 0 | 1.03 | 0.99, 1.07 | 0.12 | 0 |
| Hayley | M | 1.30 | 1.24, 1.37 | <0.001 | + | 1.36 | 1.29, 1.42 | <0.001 | + |
|  | F | 0.96 | 0.91, 1.01 | 0.13 | 0 | 1.01 | 0.96, 1.07 | 0.67 | 0 |
| Potton | M | 1.41 | 1.36, 1.46 | <0.001 | + | 1.45 | 1.40, 1.50 | <0.001 | + |
|  | F | 0.95 | 0.90, 1.00 | 0.06 | 0 | 1.02 | 0.97, 1.06 | 0.53 | 0 |
| Waresley | M | 1.35 | 1.28, 1.42 | <0.001 | + | 1.40 | 1.34, 1.47 | <0.001 | + |
|  | F | 0.92 | 0.87, 0.97 | 0.003 | - | 0.98 | 0.93, 1.03 | 0.38 | 0 |
| Weaveley | M | 1.40 | 1.32, 1.47 | <0.001 | + | 1.44 | 1.36, 1.51 | <0.001 | + |
|  | F | 0.99 | 0.93, 1.05 | 0.83 | 0 | 1.04 | 0.99, 1.11 | 0.13 | 0 |

M, Male; F, Female; CI, 95% confidence intervals; +, positive allometry; -, negative allometry; 0, isometry.

## Results

### (1) Sex differences in the allometric relationship between head width and pronotum width

Looking at head width pooled across all the sampled woodlands, head width scaled differently with pronotum width in males and females (OLS: t_1724_ = -23.21, p < 0.001; SMA: LR_1_ = 426.7, p < 0.001; Figure 1). Subsequent analysis revealed that male head width exhibited a positive slope that was significantly steeper than 1 (i.e., it was hyperallometric: OLS: F_798_ = 716.73, p < 0.001; SMA: r_798_ = 0.79, p < 0.001; Table 1), whereas female head width either exhibited a positive slope significantly lower than 1 (i.e it was hypoallometric, OLS: F_926_ = 16.79, p < 0.001; Table 1) or had a slope of 1 (i.e. it was isometric, SMA: r_926_ = 0.04, p = 0.27; Table 1) depending on the statistical method used (OLS *vs* SMA). Furthermore, sex differences in the scaling relationship between head width and pronotum width occurred in every woodland population when each woodland was tested separately (Table 1, Figure 2). Male head width was consistently hyperallometric in every woodland, regardless of whether OLS or SMA was used for the analysis (Table 1, Figure 2). Female head width, on the other hand, was isometric in every woodland in the SMA analysis, but was either hypoallometric (Buff, Cockayne Hatley, and Waresley), or isometric (Gamlingay, Hayley, Potton, and Weaveley) in the OLS analysis (Table 1, Figure 2).

**Figure 1.**
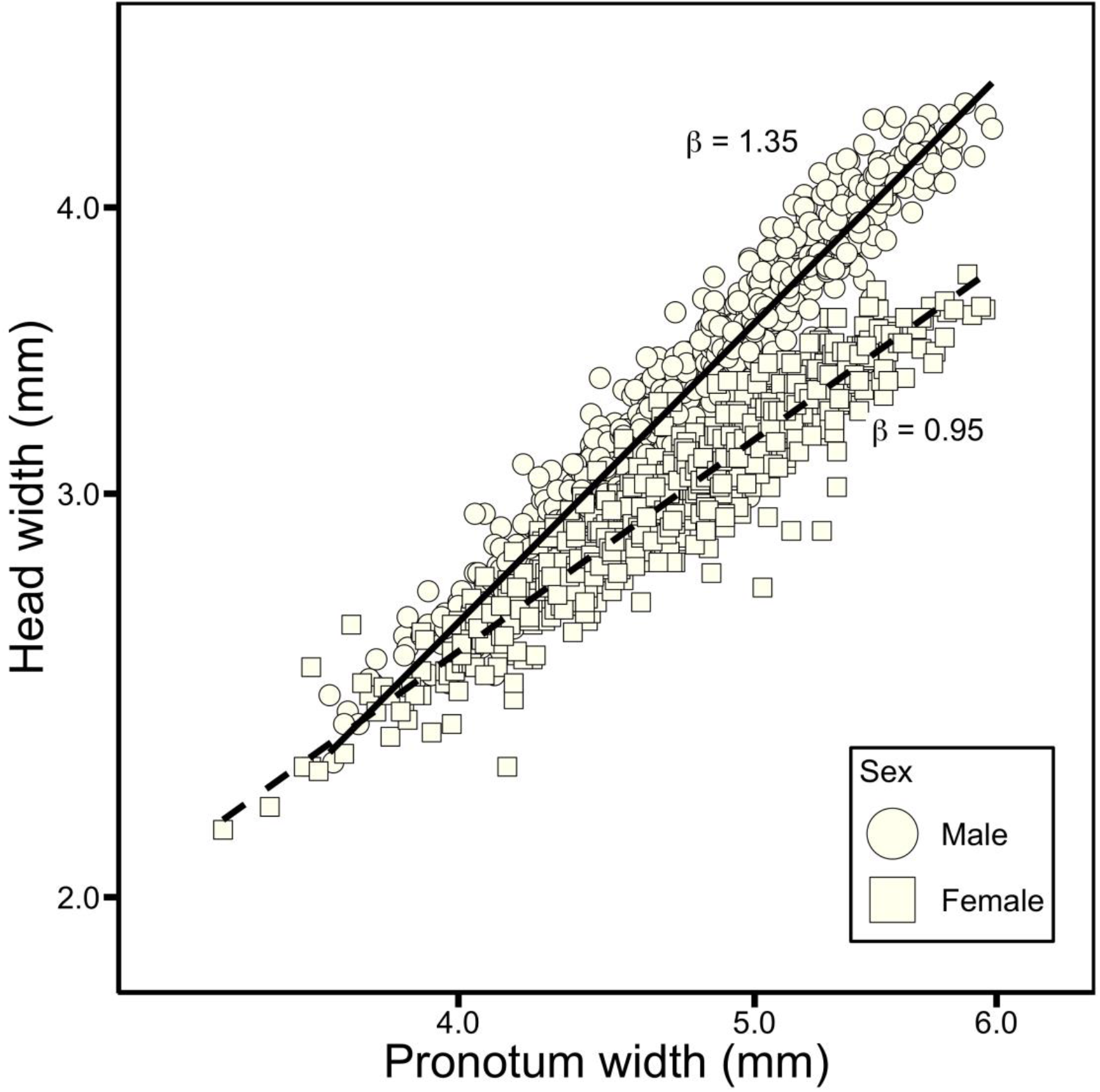
The allometric relationship between head width and pronotum width for male (circular datapoints) and female (square datapoints) *Nicrophorus vespilloides*. Lines shown are the results of ordinary least squares regressions on log-log-transformed data (solid for males, dashed for females).

**Figure 2.**
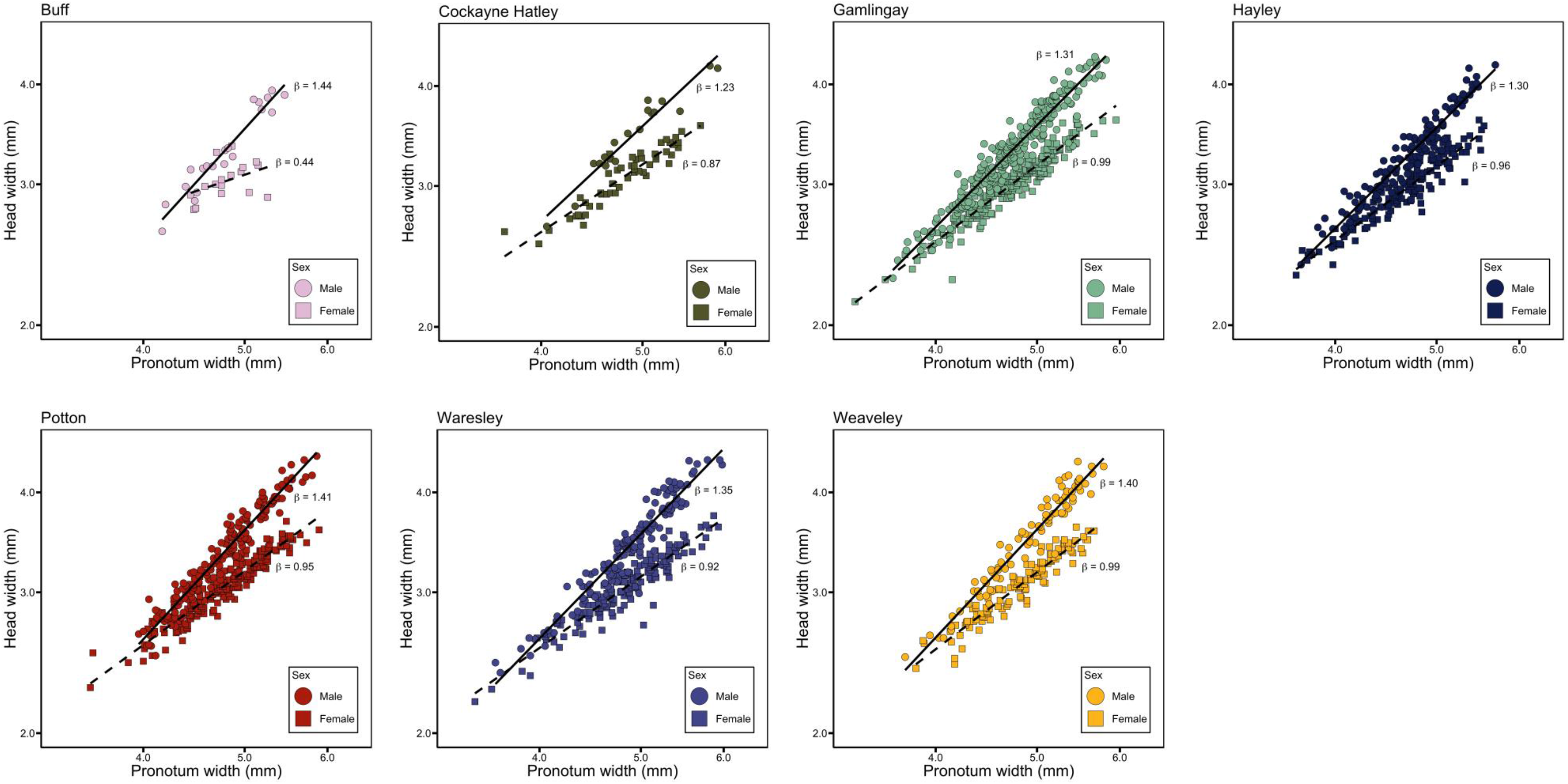
The allometric relationship between head width and pronotum width in male (circular datapoints) and female (square datapoints) *Nicrophorus vespilloides* for each woodland. Lines shown are the results of ordinary least squares regressions on log-log-transformed data.

### (2) Variation among populations in the sex difference in head size allometry

Next, we tested whether sex differences in the scaling relationship between head width and pronotum width differed among woods, initially by comparing the scaling relationship (β) across woods separately for each sex. For males, there was a small but significant difference in allometric slope between woods in the OLS analysis (OLS: F_6,786_ = 2.12, p = 0.05), though no pairwise comparisons were significant (Supplementary / Table 2A). Likewise, no significant differences in slope could be detected with the SMA analysis (SMA: LR_6,786_ = 11.29, p = 0.08). For females we also found significant differences in allometric slope between woods in the OLS analysis (OLS: F_6,914_ = 3.57, p = 0.002), with pairwise comparisons showing that the allometry in Buff was significantly shallower than in five other woods (Supplementary / Table 2B). However, no equivalent differences between females were found using the SMA analysis (SMA: LR_6,914_ = 6.11, p = 0.41).

We did, however, find that the elevation (intercept) of the allometric relationship between head width and pronotum width differed between woods, in both types of analysis, and in both males (OLS: F_6,786_ = 2.80, p = 0.01; SMA: χ^2^ = 16.26, p = 0.01), and females (OLS: F_6,914_ = 5.45, p < 0.001; SMA: χ^2^ = 26.60, p < 0.001). However, in males, no pairwise comparisons were significant in both OLS and SMA analyses (Supplementary / Tables 3A and 4A). In females, pairwise comparisons showed that the allometric elevation of Waresley was significantly lower than Cockayne Hatley and Potton (Supplementary / Tables 3B and 4B).

### (3) Variation among populations in sexual dimorphism in head width

In a final analysis, we compared whether the difference in head width between males and females varied across the different woods (i.e., the degree of sexual dimorphism), controlling for pronotum width (which is correlated with head width, but which shows no sexual dimorphism; Supplementary / Figure 3 and Table 1). In every wood males had wider heads than females (sex: F_1,1713_ = 2985.54, p < 0.001; Table 2, Table 3) but the magnitude of the difference between the sexes in head size differed among woods (significant sex x woodland interaction: F_6,1713_ = 4.67, p < 0.001; Table 2, Figure 3, Supplementary / Table 5). Weaveley and Waresley both had a significantly greater male-female head width difference than Hayley, Gamlingay, and Potton. Furthermore, Hayley had a significantly greater difference than Potton (Supplementary / Table 5).

**Figure 3.**
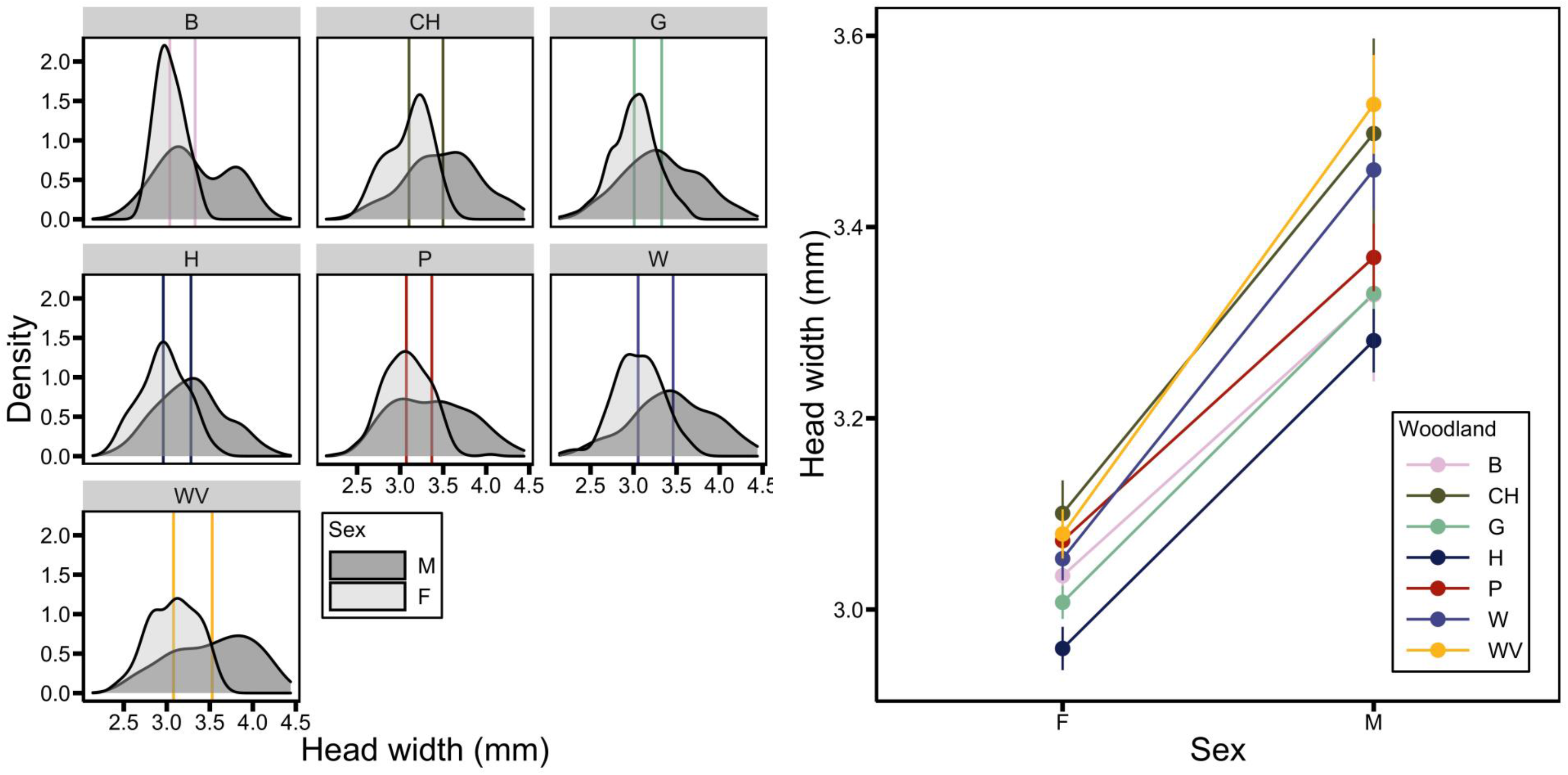
Sexual dimorphism in *N. vespilloides* head width. Left: Density distributions of head width for females (F) and males (M) across woodlands (B = Buff; CH = Cockayne Hatley; G = Gamlingay; H = Hayley; P = Potton; W = Waresley; WV = Weaveley). Vertical lines indicate median for each sex, in each woodland. Right: Plot of mean (± standard error) showing the extent of sexual dimorphism in head width across each woodland.

**Table 2.** Summary of the final ANCOVA model analysing the extent of sexual dimorphism in head width between woodland populations.

| Dependent variable | Explanatory variables | df | MS | F | p |
| --- | --- | --- | --- | --- | --- |
| head width | sex | 1 | 49.21 | 2985.54 | <0.001 |
|  | woodland | 6 | 0.93 | 56.47 | <0.001 |
|  | pronotum width | 1 | 193.02 | 11709.90 | <0.001 |
|  | sex x woodland | 6 | 0.08 | 4.67 | <0.001 |

df, degrees of freedom; MS, mean square.

**Table 3.** Quantifying woodland head width in male and female *N. vespilloides*. Mean, standard error (SE), sample size (N) and differences for head width between sexes for each woodland, with 95% confidence intervals (CI).

| Woodland | Males |  |  | Females |  |  | Difference | CI |
| --- | --- | --- | --- | --- | --- | --- | --- | --- |
|  | N | Mean | SE | N | Mean | SE |  |  |
| Buff | 20 | 3.33 | 0.09 | 17 | 3.04 | 0.04 | 0.29 | 0.08, 0.50 |
| Cockayne Hatley | 18 | 3.50 | 0.10 | 54 | 3.10 | 0.03 | 0.40 | 0.24, 0.56 |
| Gamlingay | 245 | 3.33 | 0.03 | 236 | 3.01 | 0.02 | 0.32 | 0.26, 0.38 |
| Hayley | 139 | 3.28 | 0.03 | 144 | 2.96 | 0.02 | 0.32 | 0.24, 0.40 |
| Potton | 164 | 3.37 | 0.04 | 205 | 3.07 | 0.02 | 0.30 | 0.23, 0.37 |
| Waresley | 128 | 3.46 | 0.04 | 154 | 3.05 | 0.02 | 0.41 | 0.32, 0.50 |
| Weaveley | 86 | 3.53 | 0.05 | 118 | 3.08 | 0.03 | 0.45 | 0.34, 0.56 |

## Discussion

We describe here a form of sexual size dimorphism that has previously been overlooked in the otherwise broadly monomorphic *N. vespilloides* (Al Shareefi and Cotter, 2019; Gibbs et al., 2008; Sikes et al., 2002). We show that for the same body size (pronotum width), males consistently have wider heads than females. We found the same pattern of head width dimorphism in each of the wild populations we sampled. Furthermore, for males the relationship between head width and pronotum width (body size) was hyperallometric, which is usually a characteristic of male weaponry (Emberts et al., 2021; Kodric-Brown et al., 2006). Indeed, in other insect species, head size is sometimes reckoned to be part of an individual’s armoury – under positive selection through male-male competition, and via female choice (Goyens et al., 2014; Judge and Bonanno, 2008). However, we reject that suggestion for all the reasons outlined in the Introduction. This instance of sexual size dimorphism is unlikely to be attributable to competition for mating opportunities or to mate choice.

We suggest instead that head size dimorphism is due to selection acting differently on each sex due to task specialisation during biparental care (Hedrick and Temeles, 1989; Shine, 1989; Slatkin, 1984). Previous work has shown that bite force in insects is positively associated with head size (Goyens et al., 2014) because a larger head capsule is required to accommodate the larger mandibular muscle volume that is required to close the mandibles with greater force and to chew more effectively (Bernays and Hamai, 1987; Goyens et al., 2014). Biting is central to the care-related activities in which males take the lead. Males are more likely than females to be involved in defending the nest and larvae from infanticidal takeover (Trumbo, 2007, 1994). Rival beetles bite viciously at each other’s abdomens, antennae, and legs, and consequently it is not uncommon for fights to result in injury or death (Bartlett, 1987; Müller et al., 2003; Sakaluk and Müller, 2008). A superior bite force is known to enhance performance in fights undertaken by other species (Palaoro and Peixoto, 2022). Males also take the lead when the pair converts the dead body into an edible nest for their larvae, an activity that involves scissoring off the fur or feathers with serrated mandibles and shifting, biting and shaping the carrion to round it into a ball (Eggert et al., 1998; Pukowski, 1933; Scott, 1998). Specialising in both these duties of care, therefore, demands the capacity to bite with greater force, which may have imposed selection on males to have a wider head than females. It is also possible that the females’ specialist duties of care, such as passing fluids to larvae by oral trophallaxis, require a narrower head. Perhaps this allows a better alignment of mouthparts between mother and offspring, to ensure more efficient transfer of fluids. These are hypotheses that remain to be tested in future work.

Although male head width was consistently hyperallometric, female head size was either isometric or hypoallometric, depending on whether SMA or OLS was used. These differences could simply represent a case of underestimated slopes, which is likely to occur in OLS when measurement error is present, for example, when sample sizes are low (Kilmer and Rodríguez, 2017). This may explain the marked differences between SMA and OLS. Regardless, there were obvious sex differences in the scaling relationship between head size and body size which occurred in every woodland population. While there was little consistent evidence that the slope of the scaling relationship differed among woods for either sex, we did find evidence that the elevation in the scaling relationship between head width and pronotum width varied from wood to wood. The magnitude of the difference between the sexes in head width also differed between the woods, although no such difference occurred in pronotum width (consistent with previous reports from Jarrett et al., 2017; Sun et al., 2020). This suggests that differences in pronotum width between the sexes are not driving the sex differences in head width we observe. Regarding the differences in head width between the populations, whilst it is possible that this variation is entirely environmental in origin, and due to subtle microenvironmental differences from one wood to the next in the resources available during development, no differences in the carrion resource base between these seven woods has been detected (Catherall-Ostler, 2023). It is also possible that there is additive genetic variation in head width, that is distributed differently among the woods and which persists, somehow, even though the woods are within the flight range of an individual beetle (Attisano and Kilner, 2015) and gene flow appears to be ongoing between the woods (Pascoal and Kilner, 2017; Sun et al., 2020). Genomic analyses of *N. vespilloides* from these woods has revealed that divergence at some loci is possible even when woods are less than 2km apart (Sun et al., 2020), although whether these loci are linked to the expression of head width is not yet known. Nor is it clear how such genetic divergence could persist at such a small spatial scale. It is also unclear whether the difference in head width among local populations is adaptive or not – these are all unknowns to address in future work.

Finally, we wonder whether the dimorphism in head size is confined to *N. vespilloides* or whether it will prove to be a common feature of *Nicrophorus* species. Variation among populations in the head width exhibited by another *Nicrophorus* species has already been documented in the literature. *N. microcephalus*, presumably named for its relatively small head, was once considered a separate species but is now regarded as a phenotypic variant of *N. investigator*. *N. microcephalus* is described in some detail by Power (1865). His writing hint both at sexual dimorphism in traits associated with the head, and in variation among populations of *N. investigator*, similar to the variation among *N. vespilloides* populations shown in Figure 3b: “in the head of *N. microcephalus* there is a smaller mass of vertex behind the eyes: remarkable in the male, but manifest enough in the female also…*N. microcephalus* is, as far as I have seen, a much smaller insect” (Power 1865). Comparative studies across *Nicrophorus* species might offer a way to elucidate the function of head size, particularly if detailed observations suggest that task specialisation by each sex during biparental care differs across species. For example, we might predict head width to be more monomorphic in species where females are as engaged as males in carrion nest preparation or in defending the carrion nest and larvae from attack by rivals.

In summary, we have described sexual size dimorphism in head width which, to our knowledge, has not previously been reported in wild populations of *N. vespilloides*. The natural history of *N. vespilloides* leads us to hypothesize that sexual dimorphism could result from divergent task specialisation by each sex during biparental care, rather than from sexual selection. Whether this is indeed the case, and whether it can account for the divergence between populations in absolute head width and in the extent of sexual dimorphism that we also describe here, remain to be determined in future work.

## Acknowledgements

We thank the BCN Wildlife Trust for access to Gamlingay, Hayley and Waresley Woods; the Forestry Commission for access to Potton Wood; the Tetworth Estate for access to Weaveley Wood; and Mr Michael Astor for access to Buff and Cockayne Hatley Woods. JMS was supported by a studentship from the Biotechnology and Biological Sciences Research Council’s Doctoral Training Partnership at Cambridge University. AMC was supported by a studentship from the Natural Environment Research Council’s Earth System Sciences Doctoral Training Partnership at Cambridge University. RM was supported by a BBSRC Future Leaders Fellowship (BB/R01115X/1).

## Contribution of Authors

JMS, AMC and RMK designed the study. AMC acquired the phenotypic data, which was analysed by JMS and RM. JMS and RMK wrote the manuscript.

---

**Supplementary Figure 1.**
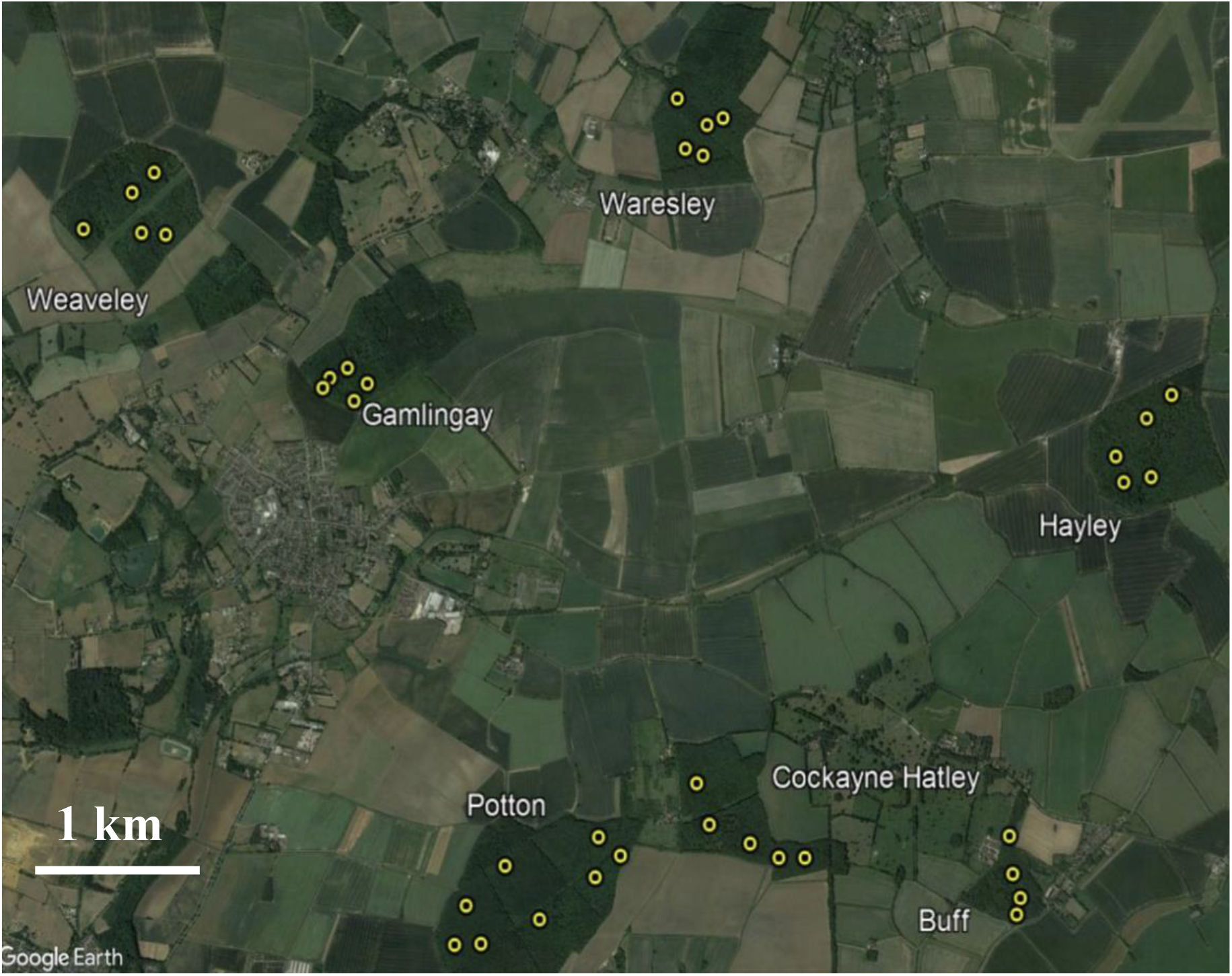
The seven woods where beetles were trapped in Cambridgeshire and Bedfordshire (UK): clockwise from top centre: Waresley Wood, Hayley Wood, Buff Wood, Cockayne Hatley Wood, Potton Wood, Gamlingay Wood, Weaveley Wood. Yellow circles depict trapping sites within each wood.

**Supplementary Figure 2.**
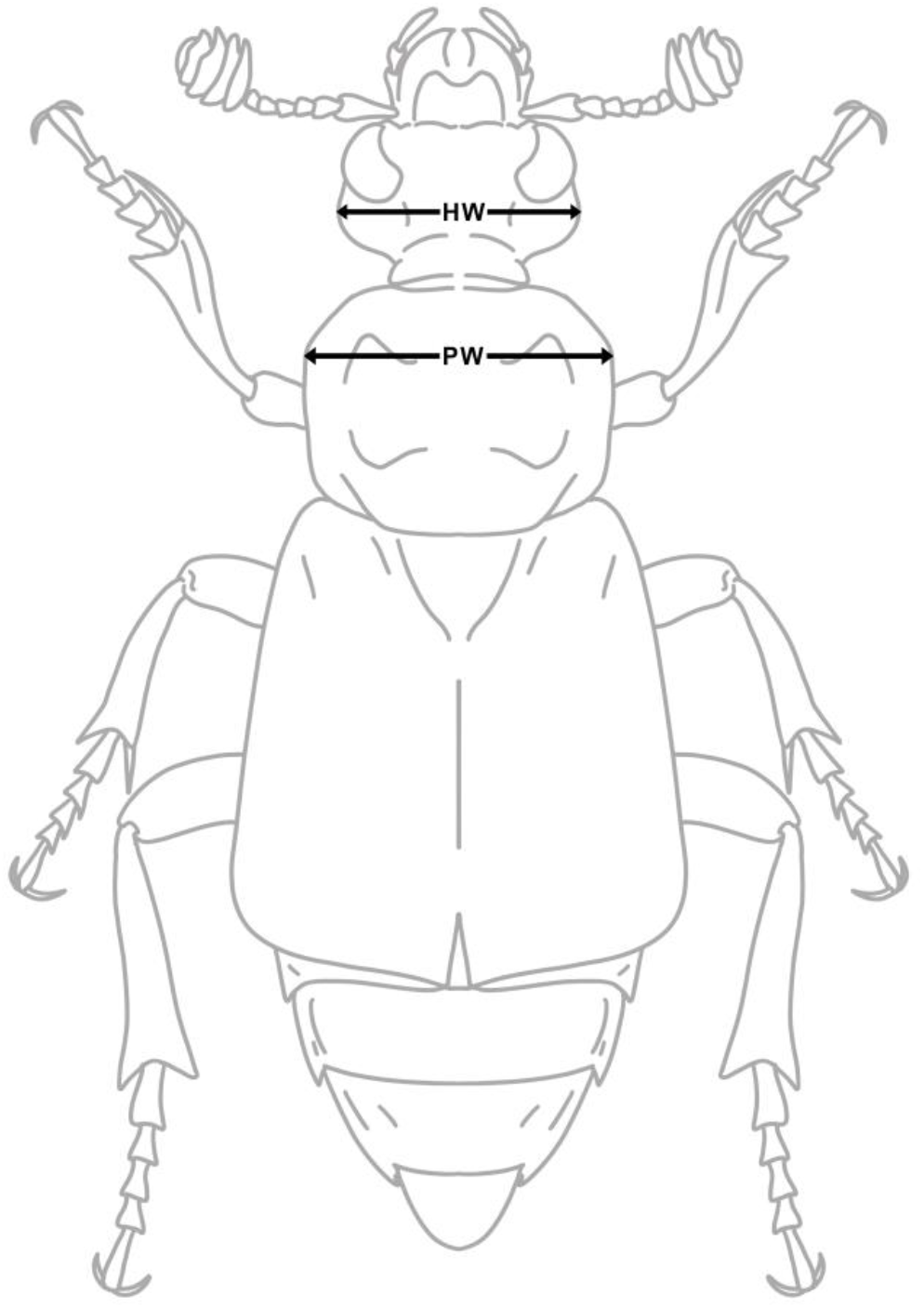
Diagram of morphological measurements (dorsal view). Head width (HW) was measured as the distance between the two widest points of the head capsule. Pronotum width (PW) was measured as the distance between the two widest points of the pronotum.

**Supplementary Figure 3.**
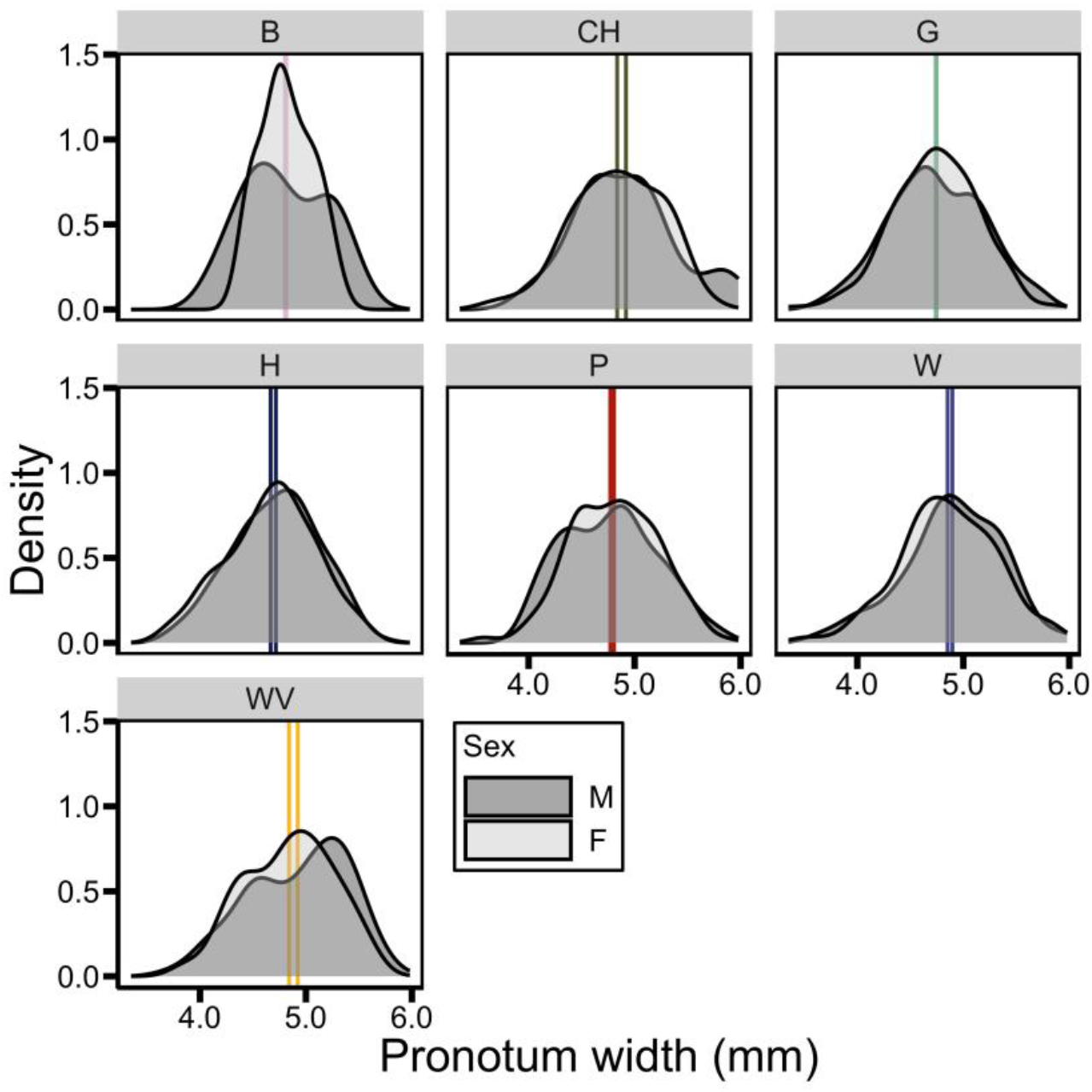
Distribution of pronotum width for males (M) and females (F) among woodlands (B = Buff; CH = Cockayne Hatley; G = Gamlingay; H = Hayley; P = Potton; W = Waresley; WV = Weaveley). Vertical lines show median for each sex, in each woodland.

**Supplementary Table 1.** Summary of the ANCOVA model analysing the extent of sexual dimorphism in pronotum width between woodland populations.

| Dependent variable | Explanatory variables | df | MS | F | p |
| --- | --- | --- | --- | --- | --- |
| pronotum width | sex | 1 | 0.08 | 0.41 | 0.52 |
|  | woodland | 6 | 1.27 | 6.77 | <0.0001 |
|  | sex x woodland | 6 | 0.11 | 0.58 | 0.75 |

df, degrees of freedom; MS, mean square.

**Supplementary Table 2.** Pairwise post-hoc comparisons using Tukey’s correction (t), testing for differences in OLS slope of head width allometry between woods (B = Buff; CH = Cockayne Hatley; G = Gamlingay; H = Hayley; P = Potton; W = Waresley; WV = Weaveley) for males (A) and females (B). p-values (p) in bold indicate comparisons with significant difference (p < 0.05).

| A) Male<br>slope<br>(OLS) | B |  | CH |  | G |  | H |  | P |  | W |  |
| --- | --- | --- | --- | --- | --- | --- | --- | --- | --- | --- | --- | --- |
|  | t | p | t | p | t | p | t | p | t | p | t | p |
| CH | 1.64 | 0.66 | - | - | - | - | - | - | - | - | - | - |
| G | 1.26 | 0.87 | -0.98 | 0.96 | - | - | - | - | - | - | - | - |
| H | 1.35 | 0.83 | -0.81 | 0.98 | 0.33 | 1.00 | - | - | - | - | - | - |
| P | 0.31 | 1.00 | -1.99 | 0.42 | -2.52 | 0.15 | -2.42 | 0.19 | - | - | - | - |
| W | 0.89 | 0.97 | -1.34 | 0.84 | -0.90 | 0.97 | -1.06 | 0.94 | 1.37 | 0.82 | - | - |
| WV | 0.43 | 1.00 | -1.79 | 0.56 | -1.81 | 0.54 | -1.86 | 0.51 | 0.26 | 1.00 | -0.93 | 0.97 |

| B) Female<br>slope<br>(OLS) | B |  | CH |  | G |  | H |  | P |  | W |  |
| --- | --- | --- | --- | --- | --- | --- | --- | --- | --- | --- | --- | --- |
|  | t | p | t | p | t | p | t | p | t | p | t | p |
| CH | -2.84 | 0.07 | - | - | - | - | - | - | - | - | - | - |
| G | -3.72 | **0.004** | -2.25 | 0.27 | - | - | - | - | - | - | - | - |
| H | -3.52 | **0.008** | -1.66 | 0.65 | 0.75 | 0.99 | - | - | - | - | - | - |
| P | -3.49 | **0.009** | -1.58 | 0.69 | 1.00 | 0.95 | 0.18 | 1.00 | - | - | - | - |
| W | -3.26 | **0.02** | -0.93 | 0.97 | 1.97 | 0.43 | 1.07 | 0.94 | 0.96 | 0.96 | - | - |
| WV | -3.73 | **0.004** | -2.21 | 0.29 | -0.21 | 1.00 | -0.83 | 0.98 | -1.03 | 0.95 | -1.85 | 0.52 |

**Supplementary Table 3.** Pairwise post-hoc comparisons using Tukey’s correction (t), testing for differences in OLS elevation of head width allometry between woods (B = Buff; CH = Cockayne Hatley; G = Gamlingay; H = Hayley; P = Potton; W = Waresley; WV = Weaveley) for males (A) and females (B). p-values (p) in bold indicate comparisons with significant difference (p < 0.05).

| A) Male elevation (OLS) | B |  | CH |  | G |  | H |  | P |  | W |  |
| --- | --- | --- | --- | --- | --- | --- | --- | --- | --- | --- | --- | --- |
|  | t | p | t | p | t | p | t | p | t | p | t | p |
| CH | -1.69 | 0.62 | - | - | - | - | - | - | - | - | - | - |
| G | -2.01 | 0.41 | 0.33 | 1.00 | - | - | - | - | - | - | - | - |
| H | -1.29 | 0.86 | 0.97 | 0.96 | 1.51 | 0.74 | - | - | - | - | - | - |
| P | -2.46 | 0.18 | -0.13 | 1.00 | -1.14 | 0.92 | -2.39 | 0.21 | - | - | - | - |
| W | -1.45 | 0.78 | 0.80 | 0.99 | 1.10 | 0.93 | -0.33 | 1.00 | 1.98 | 0.43 | - | - |
| WV | -2.82 | 0.07 | -0.58 | 1.00 | -1.84 | 0.52 | -2.85 | 0.07 | -0.88 | 0.98 | -2.53 | 0.15 |

| B) Female elevation (OLS) | B |  | CH |  | G |  | H |  | P |  | W |  |
| --- | --- | --- | --- | --- | --- | --- | --- | --- | --- | --- | --- | --- |
|  | t | p | t | p | t | p | t | p | t | p | t | p |
| CH | -2.13 | 0.34 | - | - | - | - | - | - | - | - | - | - |
| G | -0.74 | 0.99 | 2.69 | 0.10 | - | - | - | - | - | - | - | - |
| H | -0.56 | 1.00 | 2.81 | 0.08 | 0.41 | 1.00 | - | - | - | - | - | - |
| P | -1.82 | 0.53 | 0.87 | 0.98 | -2.86 | 0.07 | -2.90 | 0.06 | - | - | - | - |
| W | 0.16 | 1.00 | 4.01 | **0.001** | 2.20 | 0.30 | 1.58 | 0.69 | 4.70 | **0.0001** | - | - |
| WV | -1.14 | 0.92 | 1.81 | 0.54 | -0.97 | 0.96 | -1.22 | 0.89 | 1.42 | 0.79 | -2.76 | 0.09 |

**Supplementary Table 4.** Pairwise post-hoc comparisons using Sidak correction (t), testing for differences in SMA elevation of head width allometry between woods (B = Buff; CH = Cockayne Hatley; G = Gamlingay; H = Hayley; P = Potton; W = Waresley; WV = Weaveley) for males (A) and females (B). p-values (p) in bold indicate comparisons with significant difference (p < 0.05).

| A) Male<br>elevation<br>(SMA) | B |  | CH |  | G |  | H |  | P |  | W |  |
| --- | --- | --- | --- | --- | --- | --- | --- | --- | --- | --- | --- | --- |
|  | t | p | t | p | t | p | t | p | t | p | t | p |
| CH | 2.67 | 0.90 | - | - | - | - | - | - | - | - | - | - |
| G | 4.44 | 0.53 | 0.08 | 1.00 | - | - | - | - | - | - | - | - |
| H | 1.90 | 0.98 | 0.98 | 1.00 | 2.16 | 0.96 | - | - | - | - | - | - |
| P | 7.22 | 0.14 | 0.17 | 1.00 | 1.17 | 1.00 | 5.15 | 0.39 | - | - | - | - |
| W | 1.76 | 0.99 | 0.58 | 1.00 | 1.58 | 0.99 | 0.002 | 1.00 | 5.99 | 0.26 | - | - |
| WV | 6.97 | 0.16 | 0.30 | 1.00 | 2.24 | 0.95 | 5.90 | 0.27 | 0.06 | 1.00 | 5.69 | 0.30 |

| B) Female<br>elevation<br>(SMA) | B |  | CH |  | G |  | H |  | P |  | W |  |
| --- | --- | --- | --- | --- | --- | --- | --- | --- | --- | --- | --- | --- |
|  | t | p | t | p | T | p | t | p | t | p | t | p |
| CH | 1.57 | 0.99 | - | - | - | - | - | - | - | - | - | - |
| G | 0.24 | 1.00 | 4.97 | 0.42 | - | - | - | - | - | - | - | - |
| H | 0.20 | 1.00 | 5.40 | 0.35 | 0.004 | 1.00 | - | - | - | - | - | - |
| P | 0.98 | 1.00 | 0.55 | 1.00 | 6.64 | 0.19 | 5.25 | 0.37 | - | - | - | - |
| W | 0.009 | 1.00 | 14.26 | **0.003** | 6.66 | 0.19 | 3.77 | 0.68 | 20.7 | **0.0001** | - | - |
| WV | 0.36 | 1.00 | 2.91 | 0.86 | 0.24 | 1.00 | 0.25 | 1.00 | 2.51 | 0.92 | 7.55 | 0.12 |

**Supplementary Table 5.**
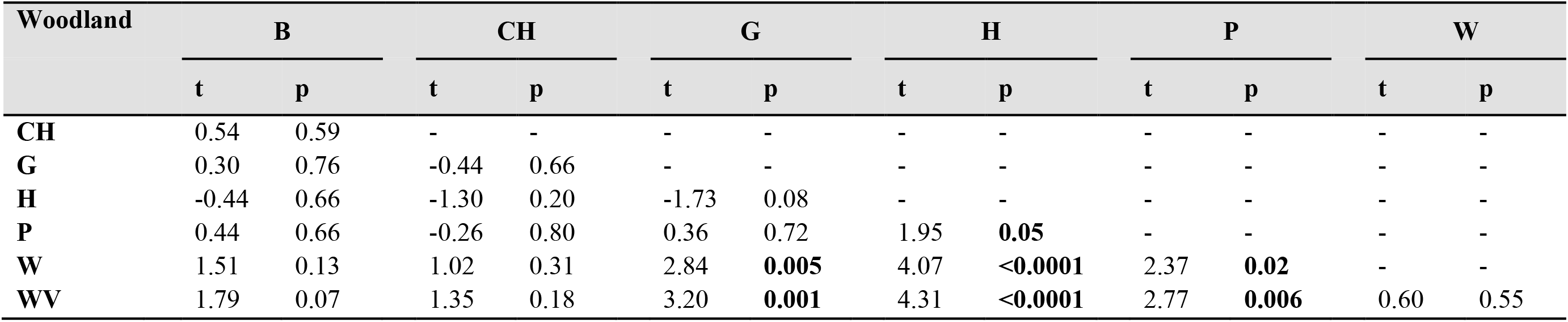
Pairwise post-hoc comparisons using Tukey’s correction (t), testing the differences in mean head width between pairs (male-female differences) between woods (B = Buff; CH = Cockayne Hatley; G = Gamlingay; H = Hayley; P = Potton; W = Waresley; WV = Weaveley). p-values (p) in bold indicate comparisons with significant difference (p < 0.05).

